# From Nuisance to Signal: Leveraging Close Relatives in Biobank-Scale Demographic Inference

**DOI:** 10.64898/2026.06.15.729614

**Authors:** Cole M. Williams, Sohini Ramachandran

**Affiliations:** Center for Computational Molecular Biology, Brown University, Providence, RI 02906; Department of Ecology, Evolution and Organismal Biology, Brown University, Providence, RI 02906

## Abstract

Biobank-scale datasets now routinely include hundreds of thousands to millions of individuals, and as sample sizes grow, close relatives become increasingly prevalent. The convention in population genetics has been to remove close relatives prior to inference, effectively treating them as a nuisance parameter. However, the consequences of this practice for demographic inference, and specifically for estimates of recent effective population size (*N_e_*), have not been rigorously evaluated. Here, we benchmark IBDNe and HapNe-IBD, two widely-used methods for inferring recent *N_e_* from identity-by-descent (IBD) segments, under a range of demographic histories and relative sampling schemes. We show that when individuals are randomly ascertained, retaining all relatives produces the least biased *N_e_* estimates; in contrast, removing even second-degree relatives inflates recent *N_e_* and induces oscillatory artifacts that “ripple”, leading to biased estimates up to ten generations into the past. We demonstrate that this ripple effect arises because close relatives contribute IBD segments that are assigned by the model to a range of ancestral ages beyond their true TMRCA, meaning their removal creates signal deficits across multiple generations simultaneously. We further show that deliberately oversampling close relatives produces severe downward bias in recent *N_e_*. To support these analyses, we develop an open-source IBD simulation pipeline using `msprime` that generates realistic IBD segments under arbitrary demographic histories and Wright-Fisher pedigrees. We provide practical guidelines for IBD simulation schemes incorporating pedigrees and argue that, in the biobank era, retaining close relatives is generally the best practice for IBD-based *N_e_* inference.

## 1 Introduction

In the past decade, biobank efforts have sequenced or genotyped millions of individuals, including the UK Biobank (500k), All of US (414k), BioVu (300k), as well as private efforts such as the Kaiser Permanente Research Bank (500k), 23andMe (14M), and Ancestry (25M) [11, 31, 3]. As datasets grow in size linearly, the number of pairs of individuals grows quadratically and so too the number of relatives, both close and distant [23]. In the UK Biobank, for instance, Nait Saada et al. [20] found that > 99% of participants have a 5th cousin or closer and almost a quarter of participants have a 2nd cousin or closer. Given a proportion of the population sampled, Shchur and Nielsen [25] provide analytical expectations of relatedness; for instance, in sampling 10% of the population (as the FinnGenn biobank has done [16], for example) half of all participants are expected to have a second degree relative (e.g., half-sibling). In this paper, we address the presence of close relatives—which we define as pairs whose time to most recent common ancestor (TMRCA) is ≤ 3—in inferring population genetic parameters, focusing on effective population size trajectory, N*_e_*(t).

Traditionally, the presence of relatives has been seen as a nuisance in population genetics, assumed to bias inference of demographic parameters, such as N*_e_*(t) and migration rates. When datasets were smaller, relatives (particularly close relatives) were rarer, and so this problem could be mitigated by removing one individual of a relative pair, resulting in a slightly smaller sample for downstream analyses. As biobanks sample higher proportions of the population of interest, this strategy will exclude large portions of biobank datasets. For example, if 10% of a population were sampled, nearly half of the dataset would need to be excluded in order to remove 3rd degree relatives (e.g., first cousins) or closer.

Population genetics has found ways to leverage distant relatedness for demographic inference, yet the consequences of continuing the practice of removing close relatives from demographic inference is unknown. For instance, the methods IBDNe [6] and HapNe-IBD [9], use identity-by-descent (IBD) segments shared between distant relatives to estimate recent N*_e_*(t). However, by default IBDNe ignores the equivalent of 2nd degree relatives; other papers utilizing IBDNe have removed other close relatives as well [27]. In the IBDNe manual, the authors recommend removing 2nd degree relatives if the population sample is not random.

Close relatives have been leveraged to estimate population sizes for decades in the field of conservation genetics [34]. For instance, close-kin mark-recapture methods (CMKR) are used to estimate census population size; by sampling a number of individuals and observing the size of the subset that are related, the population size can be estimated (i.e., more close relatives observed means a smaller population size). This is related to the work of Shchur and Nielsen [25], who derived analytical expectations using human mating patterns for the relationship between the proportion of relatives, the sample size, and the population size. From a conservation genetics lens, [33] warn against removing siblings from N*_e_*(t) estimators. The authors argue that removing siblings may be necessary to reduce bias in N*_e_*(t) estimators, but that it is virtually impossible to *know* how many siblings to remove in order to reduce the bias. Indeed, knowing how many siblings to remove to reduce bias would likely require knowing N*_e_*(t) to begin with—a circular problem. A key issue is that any attempt to resolve this circularity requires setting a hard kinship cutoff [26], yet this cutoff is inherently arbitrary: the boundary between relatedness due to recent shared ancestry and relatedness reflecting population history depends on N*_e_*(t) itself, since in smaller populations distant relatives share IBD at rates resembling close relatives in larger ones. As Jewett and the 23andMe Research Team [14] shows, relatedness classification and N*_e_*(t) inference are therefore mutually dependent.

As biobanks grow and recruitment becomes more representative, close relatives will increasingly arise as a natural consequence of sampling completeness rather than as a design artifact [8] and it is important to reassess the inclusion of genomes of close relatives in demographic inference. In this paper, we argue, in agreement with [33], that removing close relatives for N*_e_*(t) inference is ill-advised, unless some specific knowledge about the dataset necessitates it (for example, knowledge that the dataset was collected to be enriched with close relatives). We developed an IBD simulation pipeline using msprime [17] that produces realistic IBD segments under any demographic model and supports Wright-Fisher mating, as well as any user-supplied pedigree. We simulate several demographic histories using our pipeline and benchmark IBDNe [6] and HapNe [9]. We show that, when individuals are randomly ascertained, keeping all pairs produces the least biased estimate of N*_e_*(t), and removing close relatives can drastically increase recent (in the past 5 generations) estimates of N*_e_*. Importantly, we also find that removing close relatives produces oscillations in N*_e_*(t) that ripple even further into the past. In the biobank era, our results indicate that keeping all close relatives is likely the best practice for estimates of N*_e_*(t).

## 2 Methods

### 2.1 Simulations

The correct strategy for simulating IBD segments depends on the type and scale of the downstream analysis. We outline these strategies in Box 2.1. To aid in our analyses, we developed an IBD simulation pipeline that uses msprime and can be found at https://www.github.com/williamscole/ibd-sims. Our pipeline is a full end-to-end framework: it accepts any demographic model encodable as an msprime [17] Demography object, generates IBD segments under either the standard coalescent or a Wright-Fisher pedigree, and calls IBD via hap-ibd [4]. Post-processing is modular and extensible; built-in modules include IBDNe and HapNe-IBD, and users can implement arbitrary analyses by subclassing a lightweight PostProcessor interface.

We will briefly describe our pipeline here, and more details can be found in the Supplement (3.1). First, we generate a population pedigree using Wright-Fisher mating and then supply this pedigree to msprime (v1.4.1) using the fixed-pedigree mode for each chromosome. Our Wright-Fisher pedigrees only go back 25 generations, at which point the simulation switches to the standard coalescent, a hybrid approach recommended by Nelson et al. [22] (see 2.1). Our simulation pipeline supports both monogamous and non-monogamous mating, although for the purpose of N*_e_*(t) inference, the mating system should have negligible effects [32]. Indeed, we find no qualitative difference in N*_e_*(t) inference comparing monogamous and non-monogamous mating (Figure 3).

We simulated 30 chromosomes of 100 cM each with a constant recombination rate of 10^−8^. The output of msprime are tree sequences, which we could use directly to call IBD. Instead, we generated realistic genotype data (mutation rate 10^−8^, realistic minor allele frequency spectra and SNP density) and called IBD using hap-ibd [5]. Exact details for how we generated synthetic genotype data can be found in the Supplement 3.1.3.

We performed simulations on four different demographic scenarios: constant-10k, constant-100k, and an out-of-Africa 2-population (OOA2) (European and African) model from [10, 28, 1] in which we subset our analyses to the African individuals. We also used tree sequences from simulations [2] that were performed through the BALSAC genealogy of Quebec, a quasi-complete genealogy of the French Quebec [18, 29]. The founding of Quebec is considered a founder event of approximately 10,000 [29]—a demographic history that is reflected in both genetic data and the simulations through the genealogy [2]. See Supplement 3.1.2 for more details.

### 2.2 Demographic inference

We used IBDNe [6] and HapNe-IBD [9] to compare estimates of N*_e_*(t). Both methods operate on the marginal IBD segment length distribution, but HapNe-IBD boasts improvements over IBDNe: IBDNe, as noted in its original publication, suffers from oscillatory behavior. HapNe-IBD implements a regularization scheme to smooth artefactual oscillations. For each simulation, we simulate IBD between 1000 individuals, and we run these methods on downsampled IBD segment data from a subset of 250 individuals (31,125 pairs). We have three different sampling schemes for this downsampling: “oversample” (the 250 individuals are more related than a random 250 individuals), “undersample” (the 250 individuals are less related than a random 250 individuals), and “random”, where the 250 individuals are randomly sampled.

Here, oversampling represents a family-based or community-based study design, in which participants were recruited through family units or from geographically isolated communities, resulting in a dataset enriched for close relatives relative to a random draw from the population. Such designs are common in research on founder populations and rare disease genetics [24, 21], and are precisely the contexts in which IBD-based demographic inference is most frequently applied, making the effect of relative enrichment on N*_e_*(t) estimates especially practically relevant.

Detailed methodology for our sampling schemes can be found in the Supplement 3.1.4. An overview of the demographies and filtering steps can be found in Table 1.

**Table 1:** IBD Segment Distribution and Relatedness by Demographic Model.

| Category | Metric | Constant<br>($N_e = 10,000$) | Constant<br>($N_e = 100,000$) | Out-of-Africa | Quebec |
| --- | --- | --- | --- | --- | --- |
| **Random** | # of IBD segs | 251866 | 23756 | 22163 | 26046 |
|  | Pairwise IBD (cM) | 31.27 | 5.65 | 4.11 | 12.83 |
|  | Frac. pairs sharing IBD | 0.9996 | 0.5255 | 0.5073 | 0.4517 |
| **Related** | # of IBD segs | 257714 | 24722 | 22365 | 27160 |
|  | Pairwise IBD (cM) | 38.12 | 7.62 | 4.56 | 13.94 |
|  | Frac. pairs sharing IBD | 0.9996 | 0.5259 | 0.5072 | 0.4559 |
| **Unrelated** | # of IBD segs | 251100 | 23620 | 22087 | 26771 |
|  | Pairwise IBD (cM) | 30.51 | 5.53 | 4.06 | 13.06 |
|  | Frac. pairs sharing IBD | 0.9996 | 0.5242 | 0.5061 | 0.4554 |

#### Box 2.1: Simulating Identity-by-Descent Segments

There are two main categories of simulators that generate IBD segments: **demographic simulators** and **pedigree simulators**.

##### Demographic simulators

These include population software packages like msprime [17] and SLiM [13, 12]. Here we will focus on msprime, which we used for our simulations.

###### Standard coalescent

fast and memory efficient. Useful for population-level analyses, but not suitable when pairwise IBD structure is analyzed or assumed. In large cohorts, the coalescent allows individuals to have more simulated ancestors than are biologically possible [22]. Whereas a diploid individual has at most 2*^t^* distinct ancestors at generation t, the coalescent treats lineages as coalescing independently, implicitly exceeding this constraint. As a result, recent ancestry is spread too uniformly across pairs rather than being concentrated into a smaller number of genuinely closely related individuals, producing too few pairs sharing the large, genome-wide correlated IBD that characterizes true close relatives.

###### Discrete-time Wright Fisher (DTWF)

slower, but creates more realistic pairwise IBD structure, particularly for close relatives who are expected to share IBD across multiple chromosomes simultaneously. To balance accuracy and computational efficiency, [22] suggest a hybrid approach: use DTWF for the recent past and revert to the standard coalescent beyond a switchover point motivated by the minimum IBD segment length of interest in downstream analyses—segments originating deeper in the past will be too short to detect regardless of the simulation model, so the coalescent approximation is harmless there. A conservative buffer beyond this switchover is advisable to account for variance in segment lengths around their expectation.

Currently, DTWF can be specified in msprime, but with an important caveat: if each chromosome is simulated separately, a *new* population pedigree will be created for each chromosome, breaking cross-chromosome correlations within pairs. In msprime, the solution is to simulate a single genome-length contig with a 0.5 recombination rate delineating chromosomes.

For analyses operating on the marginal IBD segment length distribution (such as IBDNe), DTWF with chromosomes simulated separately will produce, in expectation, the same total IBD and segment length distribution as the single genome-length contig strategy. The difference is in the pairwise structure: the former diffuses IBD across more pairs, while the latter correctly concentrates IBD into close relative pairs. This distinction is consequential for analyses that involve identifying or filtering close relatives, where it is critical that all chromosomes follow the same shared population pedigree, so that a pair’s relatedness is expressed consistently genome-wide rather than independently on each chromosome.

###### Fixed-pedigree

msprime also allows users to simulate coalescence through a fixed pedigree. A benefit of this is that chromosomes can be simulated independently, since each chromosome traces through the same shared population pedigree. A drawback is that lineages may reach pedigree founders at different generations. Those reaching founders early are held while remaining lineages continue coalescing and accumulating recombination events, creating an asymmetric time jump in which early-founder lineages rejoin a genealogy that has already evolved for additional generations, resulting in systematically different recombination histories than they should have.

###### Our simulation pipeline

we use msprime’s fixed-pedigree mode in our pipeline for Wright-Fisher (WF) simulations. We first generated a fixed WF pedigree given the demography and then provide the fixed pedigree to msprime so that chromosomes can be simulated separately, improving computation time. Our pipeline also allows for standard coalescent and other user-provided pedigrees.

Regardless of the exact simulation scheme, both SLiM and msprime output tree sequences, from which IBD can be called directly. Genotype/sequence data can also be created from the tree sequences, and IBD can be called on that as well. We have called IBD in both ways, and found a critical difference between the two methods. IBD called from the tree sequences will be segmented as the tree topology changes, even if the node from which that IBD segment descended stays the same. IBD called from a VCF generated from the tree sequence would not split the IBD segment. Thus, we recommend merging IBD segments called directly from tree sequences that (1) end/start at the same position and (2) share the same ancestor node.

##### Pedigree simulators

Pedigree simulators simulate specific pedigree relationships and the IBD shared between them. We will focus our discussion on ped-sim [7], which incorporates both sex-specific recombination maps and a crossover interference model, creating realistic IBD segments between close relatives. ped-sim can simulate arbitrarily distant relatives, but the benefits of sex-specific recombination and the interference model are negligible for distant relatives. Additionally, distant relatives often do not share IBD [35], so many iterations of the simulation must be done to achieve even a small number of IBD-sharing distant relatives. Jewett and the 23andMe Research Team [15] present a method for efficiently simulating IBD-sharing distant relatives.ped-sim outputs exact IBD segments, which represent a clean ground truth for pedigree IBD, but with the implicit assumption that founders are unrelated. This is a meaningful truncation: IBD from deep demographic history is excluded by construction. ped-sim also allows an input VCF to generate sequence data for simulated relatives from which IBD can be called. However, this fundamentally changes what is being measured: called IBD will be a mixture of pedigree IBD and background IBD shared between founders, where the latter reflects a latent demographic history and any sampling biases present in the input VCF. Unlike exact IBD, this background component cannot be decomposed or attributed to specific ancestors. The choice between exact and called IBD in ped-sim therefore depends on whether the goal is a clean pedigree ground truth or a more realistic mimicry of real data—but the latter comes at the cost of interpretability.

###### Takeaway

the choice of simulator and simulation scheme depends on the downstream analyses. For realistic IBD sharing between close relatives, pedigree simulators such as ped-sim are the best option. For simulations requiring a specific demographic history, msprime or SLiM are required. For analyses that operate on the marginal IBD segment length distribution, such as the Ne inference step of IBDNe, standard coalescent simulations are sufficient, since the expected length distribution is preserved across simulation schemes. For analyses in which pairwise IBD is examined, such as relative detection or the relative-filtering step that precedes demographic inference, a shared-pedigree WF simulation is necessary to correctly reproduce the concentration of IBD into close relative pairs.

## 3 Results

N*_e_*(t) estimates for simulated demographic histories.

Our N*_e_*(t) benchmarking results are shown in Figure 1. The IBDNe results (left) panel cleanly demonstrate the effect of relatives in estimation of recent N*_e_*(t). Across all demographic histories, the “unrelated” IBD input results in an overestimation of recent N*_e_*(t) and the “related” IBD input drastically underestimates recent N*_e_*(t). The “random” IBD input, which randomly samples relatedness from the population, tracks the true N*_e_*(t) well in both the recent and more distant past. For HapNe-IBD, the effect of filtering of close relatives on N*_e_*(t) is attenuated—perhaps by the same smoothing process that reduces the oscillatory nature of the inferred N*_e_*(t).

**Figure 1:**
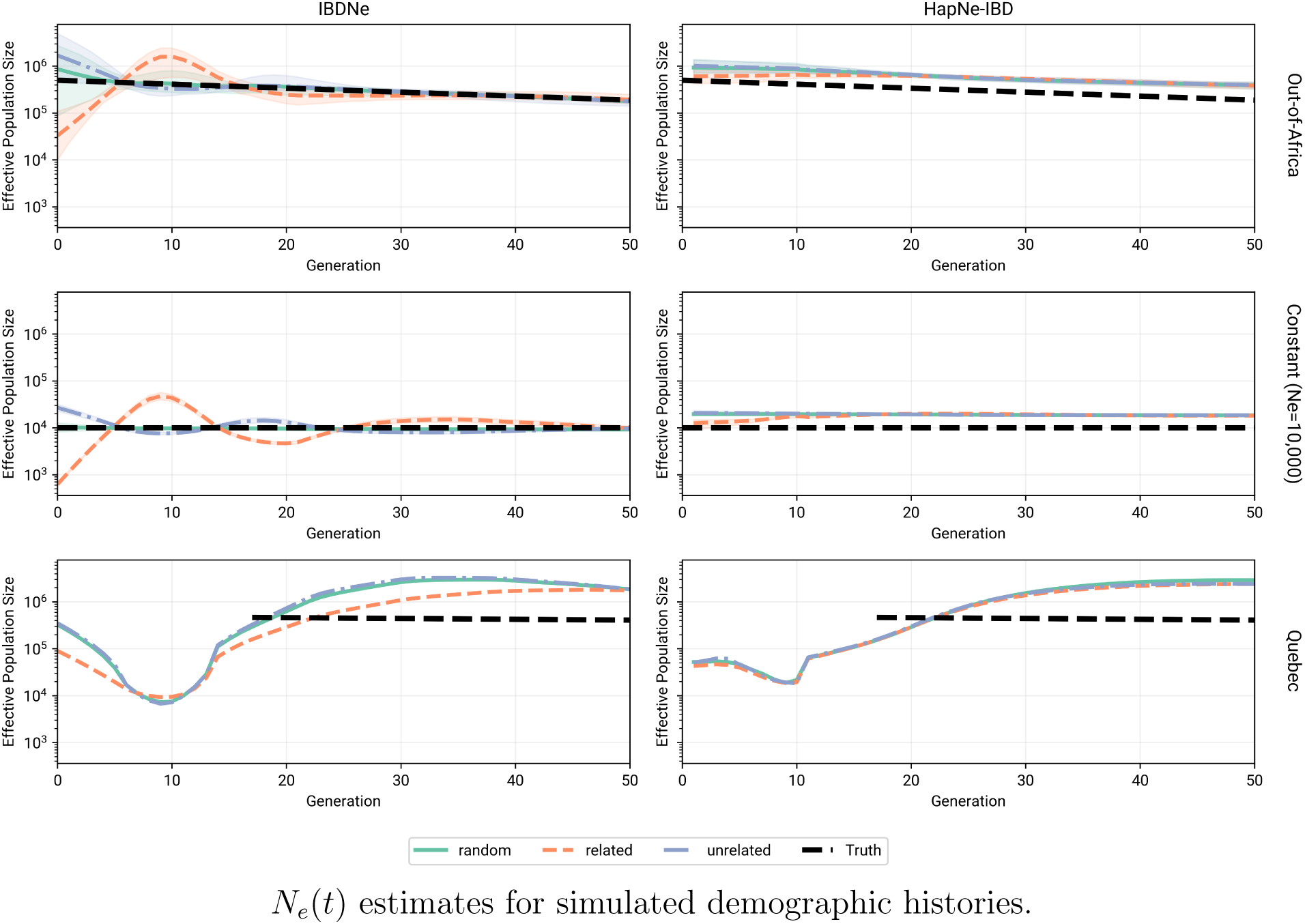
For each demographic history, 25 instantiations of the simulation was run, resulting in 25 IBD datasets for each simulation. For each IBD dataset, we filtered the input data for IBDNe [6] and HapNe-IBD [9]: related (oversampled relatedness), unrelated (undersampled relatedness), and random filtering. The mean N*_e_*(t) across the 25 IBD datasets is shown in the figure with 95% confidence interval shaded. The true N*_e_*(t) is shown as the black dashed line. For the Quebec simulations, N*_e_*(t) starts at g = 18 generations ago, as this is the end of the genealogy where the simulation reverts to the same OOA2-African demographic history.

We computed the root mean squared error (RMSE) of log_10_ N*_e_* estimates against truth for both methods across two time scales: the recent past (0–10 generations) and the more distant past (0–50 generations) (Table 2). Generally, IBDNe achieves lower RMSE than HapNe-IBD across both time scales and demographic scenarios under random sampling, though IBDNe’s advantage is concentrated in the more distant past. HapNe-IBD’s regularization produces more stable estimates in the very recent past at the cost of overall accuracy. Notably, IBDNe’s built-in relative filtering (filtersamples=True) only improves estimates when relatives are deliberately oversampled; under random or undersampled conditions it increases RMSE, consistent with our recommendation that relative filtering should be avoided unless there is clear evidence of recruitment bias. Quebec RMSE is notably higher than for other demographic scenarios, for two reasons. First, RMSE is only evaluated against known N*_e_*(t) starting at g = 18, when the simulation transitions to an Out-of-Africa African demographic history. Second, most IBD segments in the Quebec simulation originate from within the BALSAC genealogy rather than the Out-of-Africa portion, leaving both methods without sufficient coalescent signal to accurately track N*_e_*(t) at g < 18, resulting in upward-biased estimates and elevated RMSE.

**Table 2:** RMSE of log_10_ N*_e_* Ne estimates vs. truth.

| Demographic scenario | Filtering | IBDNe<br>(0–50 gen) | IBDNe<br>(0–10 gen) | HapNe-IBD<br>(0–50 gen) | HapNe-IBD<br>(0–10 gen) |
| --- | --- | --- | --- | --- | --- |
| Out-of-Africa | random | 0.126 | 0.233 | 0.287 | 0.292 |
|  | random<br>(filtersamples=True) | 0.104 | 0.188 |  |  |
|  | related | 0.339 | 0.668 | 0.258 | 0.149 |
|  | related<br>(filtersamples=True) | 0.093 | 0.175 |  |  |
|  | unrelated | 0.119 | 0.224 | 0.297 | 0.319 |
|  | unrelated<br>(filtersamples=True) | 0.119 | 0.224 |  |  |
| Constant<br>$N_e = 100,000$ | random | 0.073 | 0.120 | 0.295 | 0.298 |
|  | random<br>(filtersamples=True) | 0.107 | 0.184 |  |  |
|  | related | 0.365 | 0.699 | 0.273 | 0.142 |
|  | related<br>(filtersamples=True) | 0.070 | 0.134 |  |  |
|  | unrelated | 0.131 | 0.239 | 0.296 | 0.296 |
|  | unrelated<br>(filtersamples=True) | 0.131 | 0.239 |  |  |
| Constant<br>$N_e = 10,000$ | random | 0.031 | 0.030 | 0.281 | 0.292 |
|  | random<br>(filtersamples=True) | 0.102 | 0.170 |  |  |
|  | related | 0.342 | 0.641 | 0.259 | 0.172 |
|  | related<br>(filtersamples=True) | 0.104 | 0.172 |  |  |
|  | unrelated | 0.125 | 0.215 | 0.286 | 0.312 |
|  | unrelated<br>(filtersamples=True) | 0.125 | 0.215 |  |  |
| Quebec | random | 0.880 | 1.236 | 0.769 | 1.135 |
|  | random<br>(filtersamples=True) | 0.853 | 1.250 |  |  |
|  | related | 0.851 | 1.369 | 0.755 | 1.189 |
|  | related<br>(filtersamples=True) | 0.860 | 1.279 |  |  |
|  | unrelated | 0.890 | 1.237 | 0.746 | 1.141 |
|  | unrelated<br>(filtersamples=True) | 0.890 | 1.237 |  |  |

In the Out-of-Africa African simulations, there is a slight overestimation of recent N*_e_*(t) even when close relatives are randomly sampled. We believe that this is because our sample (n = 250) is small relative to the population size (approximately 425k), and it is unlikely, in any given simulation, for close relatives to be present,leaving the model without sufficient recent IBD signal to accurately estimate N*_e_*(t). We see a similar result for a constant-size N*_e_* = 100, 000 (Figure S4). To investigate this, we increased the sample size from 250 to 1000—representing a 16-fold increase in the number of pairs—and found that RMSE decreased substantially for IBDNe in the large N*_e_* simulations (Out-of-Africa and N*_e_* = 100, 000; Table 3); for instance, the RMSE of log_10_ N*_e_* decreased from 0.233 to 0.066 for IBDNe in the Out-of-Africa simulation. RMSE did not meaningfully decrease for HapNe-IBD, suggesting that its regularization procedure, rather than signal sparsity, is the dominant source of error regardless of sample size. For N*_e_* = 10, 000, RMSE did not decrease despite sampling increasing from 2.5% to 10% of the population, suggesting that the method was already adequately powered at the smaller sample size. Together, these results suggest that the relevant quantity for recent N*_e_* inference is not sample size alone, but the expected number of close relative pairs in the sample, ∼ n^2^/N*_e_*.

**Table 3:** RMSE of log_10_ N*_e_* Ne estimates vs. truth by sample size (random node sampling only)

| Demographic scenario | $n$ | IBDNe<br>(0–50 gen) | IBDNe<br>(0–10 gen) | HapNe-IBD<br>(0–50 gen) | HapNe-IBD<br>(0–10 gen) |
| --- | --- | --- | --- | --- | --- |
| Out-of-Africa | 250 | 0.126 | 0.233 | 0.287 | 0.292 |
|  | 1,000 | 0.045 | 0.066 | 0.268 | 0.268 |
| Constant $N_e = 10,000$ | 250 | 0.031 | 0.030 | 0.281 | 0.292 |
|  | 1,000 | 0.035 | 0.036 | 0.281 | 0.293 |
| Constant $N_e = 100,000$ | 250 | 0.073 | 0.120 | 0.295 | 0.298 |
|  | 1,000 | 0.018 | 0.024 | 0.295 | 0.297 |

Next, we investigated the effect of filtering out close relatives on the marginal IBD distribution, focusing on IBDNe. To do this, we used the following equation from Browning and Browning [6] to compute the most likely age of an IBD segment given its length and the population trajectory:

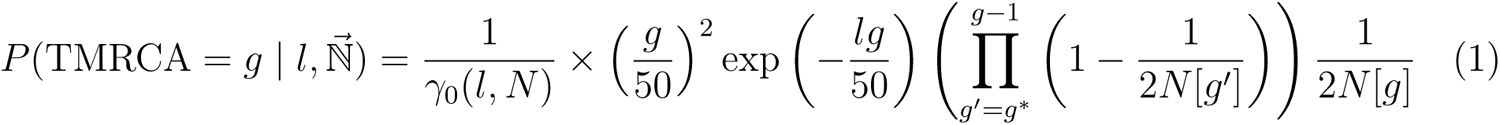

Note that some of the conditional terms from [6] have been omitted; e.g., this equation assumes that IBD segments are not at chromosome ends. Over a range of g = 1..100, we computed this probability for each segment length and selected g that maximized the probability as the most likely age of the IBD segment given the population size trajectory. Thus, for each segment we calculated: (1) the TMRCA of the pair (which is not necessarily the age of the IBD segment), and (2) the most likely age of the IBD segment, given the population size trajectory. In Figure 2, we show, for each set of segments whose most likely age is g, which set of relatives (as defined by their TMRCA) contributed those IBD segments in the Out-of-Africa African model. Figure 2 can be used to visualize the information lost by excluding relatives. TMRCA g = 3 relatives, for example, contribute 10% of the IBD segments whose most likely age is g = 1, and by removing 1st-4th degree relatives, 97% of the coalescent signal is lost for g = 1. The removal of these close relatives has far-reaching effects: 1st-4th degree relatives contribute 17% of the coalescent signal at g = 8, demonstrating the ripple effect of removing close relatives.

**Figure 2:**
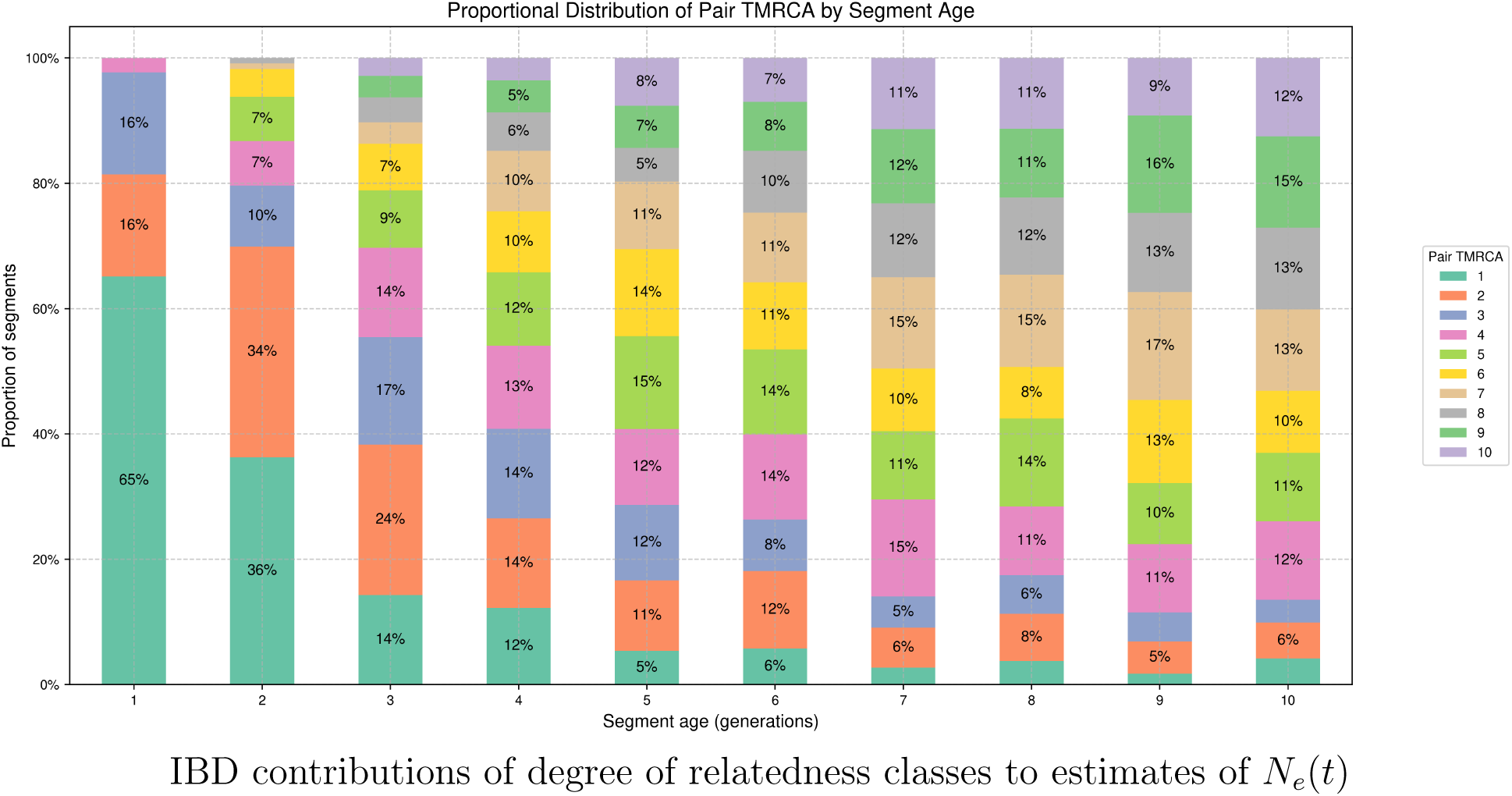
A breakdown of IBD segments, grouped by their most likely segment age (x-axis) given an OOA2-African population size trajectory. Each bar is broken down by the TMRCA of the pair (roughly equivalent to the degree of relatedness) that contributed the IBD segments that fall into that age bin. For example, of IBD segments whose most likely age is g = 1, 65% of those segments come from a pair whose TMRCA is g = 1, whereas 16% of those segments come from a pair whose TMRCA is g = 2. Close relatives (low TMRCA) contribute less to segment age bins whose most likely age is older, but the contribution is non-zero (e.g., 6% of segments whose most likely age is g = 6 generations come from half-siblings (TMRCA=1).

## Discussion

The unique, realized history of a population—the pedigree structure through which mutations arise and percolate to descendants via Mendelian inheritance—shapes genetic variation in ways that classic, population genetic models average over (e.g., Wakeley et al. [30], which established that a fixed population pedigree alters the distribution of recent coalescence times). Understanding when, how, and why this occurs requires integrating explicit genealogical structures into demographic models and data analysis. Here, we illustrate this dynamic by evaluating the estimation of recent effective population size over time, N*_e_*(t), from identity-by-descent (IBD) segment distributions (Figure 1). Our results demonstrate that the conventional practice of systematically discarding close relatives introduces severe ascertainment bias, overestimating recent population sizes and causing a distortion that ripples into the more distant past (Figure 1; see also Browning and Browning [6] and Fournier et al. [9]). To help researchers navigate these complexities, we provide specific guidelines for simulating genotype data and IBD segments within pedigree frameworks (Box 1), for future studies to evaluate how life histories and population-scale events fundamentally influence genetic variation.

Close relatives contribute IBD segments spanning a wide range of lengths, from very short (2-5 cM) to very long (50+ cM). Because IBDNe and HapNe-IBD operate on the marginal segment length distribution, shorter segments from close relative pairs are assigned older inferred TMRCAs than the pair’s true coalescence time (Figure 2). We show that this underlies the ripple effect: removing close relatives creates signal deficits across multiple inferred age bins simultaneously, distorting N*_e_*(t) estimates up to ten generations into the past. That said, the overall IBD segment length spectrum is dominated by short, old segments, few of which are contributed by close relatives, meaning their removal has negligible impact on inference in the more distant past. We therefore do not believe prior analyses using IBDNe are invalidated by our findings; N*_e_*(t) estimates do converge on the truth beyond the first several generations, and most published analyses already treat very recent estimates with caution.

A key point we wish to make is that close relatives contain signals of demographic history, not simply noise. The frequency of close relatives in a sample is itself a product of demographic history; in smaller or more isolated populations, close relatives are more prevalent, and their IBD sharing patterns directly reflect recent N*_e_*(t). Discarding close relatives implicitly assumes that only distant relative pairs carry information about demographic history. Indeed, our results show that retaining close relatives improves N*_e_*(t) estimation, not because the methods we compare explicitly model them, but because IBD segments from close relatives contribute legitimate coalescent signals that the marginal length distribution correctly absorbs. The decision to remove relatives also introduces a circularity that is rarely acknowledged: setting a principled kinship cutoff requires knowing N*_e_*(t) to begin with, since the boundary between recent shared ancestry and population-level relatedness shifts with demographic history [14]. Unless a researcher has clear evidence of recruitment bias—for instance, a family-based cohort design in which relatives were deliberately enrolled—we recommend against removing relatives prior to IBD-based N*_e_*(t) inference. As biobanks grow, if recruitment becomes representative, the frequency of close relatives in a sample will increasingly reflect true population structure rather than sampling artifacts, making their retention not just defensible but desirable. Indeed, recent recommendations for future biobank design suggest that sampling high proportions of a population such that relatives arise naturally is preferable to explicit family-based recruitment [8], a design principle that our results support from a demographic inference perspective.

Several limitations of our study warrant mention. Our simulated genotype data contained no sequencing or phasing errors, representing an idealized scenario in which IBD detection is more accurate than in practice; in real data, IBD segments should be merged post-hoc to correct for splits introduced by marker gaps, genotyping error, and phasing artifacts. Finally, our non-overlapping generation model with non-monogamous mating produces only 2nd degree (half-sibling) and 4th degree (half-cousin) relative classes, with 3rd degree relatives essentially absent; real datasets contain a continuous spectrum of relatedness degrees, and the effect of filtering may differ when avuncular and grandparent-grandchild pairs are present.

Ultimately, there remains an underappreciated distinction between genealogical and genetic relatedness—a complex relationship often ignored in both theoretical population genetic models and state-of-the-art inference approaches. As biobank datasets expand to include millions of individuals, we urge a methodological shift away from treating relatives as a nuisance parameter to be discarded wholesale, and instead suggest leveraging these relative pairs to study the rich interplay between a population’s ecology, evolution, and its pedigree.

## Acknowledgements

This work was supported by National Institutes of Health T32 GM128596 (C.M.W. and S.R.), R35 GM139628 (S.R.), and National Science Foundation GRFP (C.M.W.). Analyses were conducted using computational resources and services at the Center for Computation and Visualization, Brown University.

## Supplementary Material

### 3.1 Methods

#### 3.1.1 Creating Wright-Fisher Genealogies

Let N*_e_*[g] be the effective population size at generation g. We assume an equal number of males and females, such that each individual in generation g−1 picks a mother and father from N*_e_*[g]/2 individuals. We use msprime’s DemographyDebugger.population_size_trajectory function to query N*_e_*[g], ensuring that the correct N*_e_*[g] is used. This is important for more complex demographies. Our pipeline supports both monogamous and non-monogamous models. In the monogamous model, if an individual picks a mother who already has an offspring assigned, then the father will not be randomly chosen and will be the father of that already-assigned offspring. In the non-monogamous model, mothers and fathers are chosen randomly and independently of each other.

#### 3.1.2 Simulating with msprime

We first perform the WF simulation using the WF population pedigree generated in the step above. A demographic model is not provided, as the demography is (implicitly) encoded in the population pedigree. Once the simulation ends at g = 25 generations in the past, the result is a tree sequence, ts_1_, with un-coalesced lineages. To complete the simulation, a new simulation is run with the desired demographic history, using ts_1_ as the initial state and producing ts_2_, in which all lineages are coalesced.

For the Quebec simulations, we start with ts_2_ as provided by Anderson-Trocmé et al. [2]. We refer the reader to the original paper for details on these simulations. Briefly, they used msprime’s fixed-pedigree mode to simulate through the BALSAC genealogy [18, 29] to create the equivalent of ts_1_ and then used an Out-of-Africa (European) demographic history to complete the simulation for the un-coalesced lineages in order to create ts_2_. These simulations were run on 22 autosomes with realistic human recombination maps.

#### 3.1.3 Creating realistic genotype data

We simulate genotype data using ts_2_. First, we place mutations on the tree sequences (mutation rate 10*^−^*^8^). Next, we downsample the sites in the tree sequence to achieve a marker density of approximately 600 SNPs/Mb (∼ 1.8M SNPs genome-wide) in order to accurately call small IBD segments. Browning and Browning [4] found 50% power to detect 2 cM IBD segments with 550K SNPs, so the ∼ 3x SNP density we use should be sufficient.

We sampled sites from the minor allele frequency (MAF) histogram of the UK Biobank, using the following intervals: zero (fixed), (0, 0.01], (0.01, 0.05], (0.05, 0.1], and then four intervals of 0.1 width from 0.1 to 0.5. We then sampled, from the tree sequence, bi-allelic sites from these intervals to match the desired SNP density. These genotype data were then stored in a VCF for IBD calling.

#### 3.1.4 Sampling IBD segments

For a pair of individuals (i, j), we denote the total length (in cM) of IBD segments they share S*_i,j_*. If G is the total length of the haploid genome, then 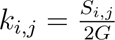 denotes the proportion of IBD sharing for the pair. Now consider t ∈ [0, 1] to be an IBD sharing proportion threshold. If k*_i,j_* > t, the pair are considered “related” and if k*_i,j_* < t, the pair are considered “unrelated”. In this study, we use t = 0.0883, which represents a lower bound separating 3rd and 4th degree relatives [19].

We create a graph G using networkx in which individual i is represented as node n*_i_*. An edge is added between nodes n*_i_* and n*_j_* if k*_i,j_* > t. Our sampling schemes operate on this graph to produce four sampling conditions, each retaining a fraction f of the full cohort of N individuals, giving us ceil(N f). We used f = 0.25, unless otherwise noted.

##### Filtering: related

To construct a subsample enriched for close relatives, we iterate through all pairs (i, j) with an edge in G in descending order of k*_i,j_*. At each step, both individuals in the pair are added to the selected set. Iteration continues until the target size is reached or all related pairs are exhausted. If the number of individuals involved in related pairs falls short of the target, the remainder is filled by drawing uniformly at random (without replacement) from the unselected pool.

##### Filtering: unrelated

To construct a subsample purged of close relatives, we seek a large independent set of G. We process each connected component of G separately. Isolated nodes (singletons) are admitted directly. For each non-trivial component, we apply a greedy algorithm: at every iteration, all candidate nodes are sorted in ascending order of their current degree within the component, and the lowest-degree node is tentatively selected. The node is added to the output set only if it shares no edge with any previously selected node; the process then repeats until no further node can be added without violating independence, or the global target is reached. If the total number of selected nodes exceeds the target, a random subset of size Nf is drawn from them.

##### Filtering: random

As a baseline condition, Nf individuals are drawn uniformly at random without replacement from the full cohort, without regard to relatedness.

### 3.2 Software

#### 3.2.1 hap-ibd [5]

hap-ibd was run with default parameters.

#### 3.2.2 IBDNe [6]

We ran IBDNe with default parameters, including a minimum IBD segment length of 2 cM. We ran it with `filtersamples=True`, which removes 2nd degree relatives, and `filtersamples=False`.

#### 3.2.3 HapNe-IBD [**9**]

HapNe-IBD was run with default parameters. We modified the HapNe-IBD code in order to support our 30 equal-length chromosome genome. The software normally takes as input the genome build, which it uses to define chromosome arms. It then computes the IBD segment length spectra for each chromosome arm. We modified the code such that it works on 60 chromosome arms, each of 50 Mb.

### 3.3 Results

**Figure 3:**
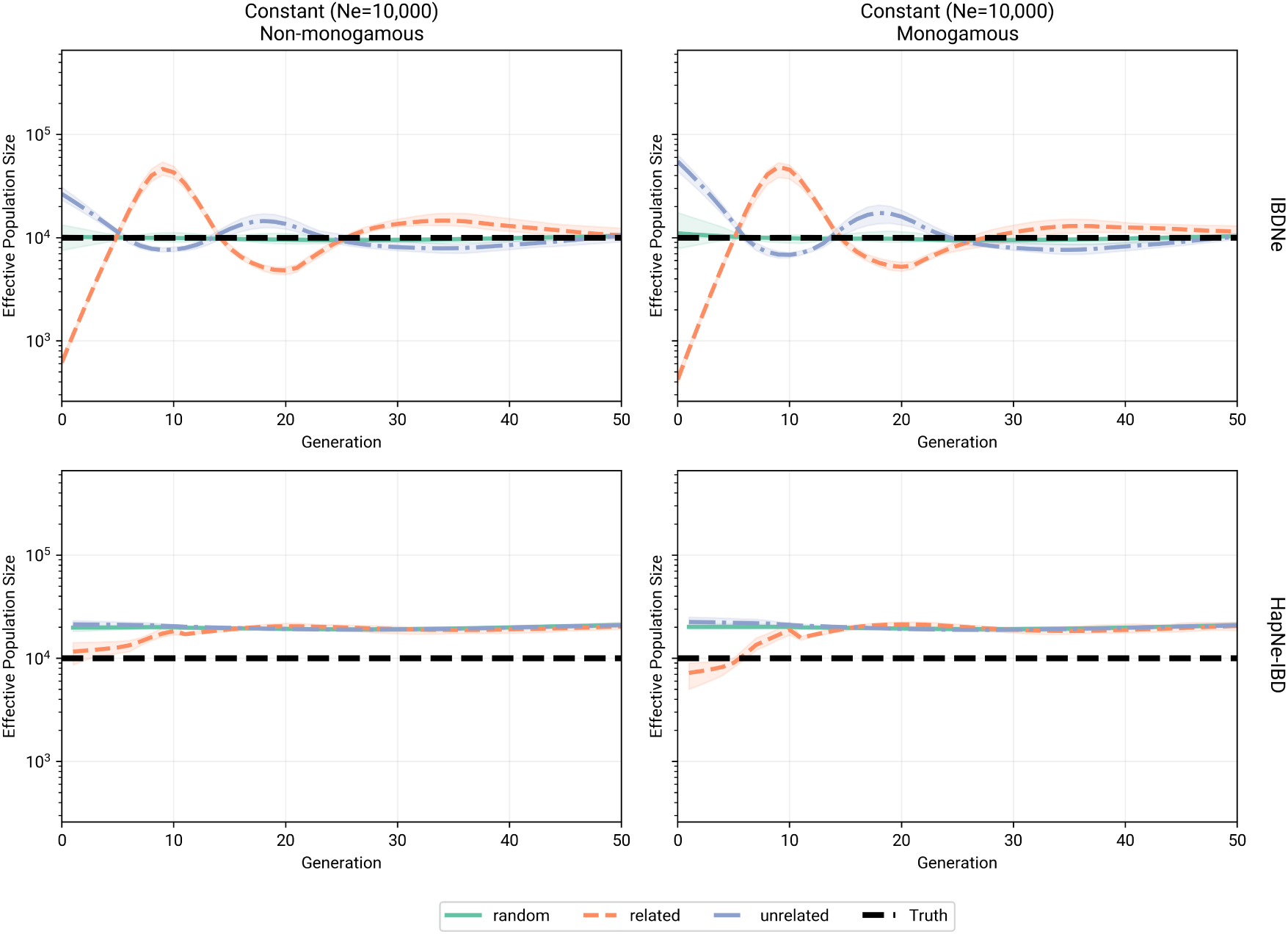
N*_e_*(t) inference for constant population size N*_e_* = 10, 000 for non-monogamous (left panels) and monogamous (right panels) mating schemes. Results for IBDNe (top panels) and HapNe-IBD (bottom panels) are shown. Qualitatively, we see no difference in N*_e_*(t) inference, in agreement with [32].

**Figure 4:**
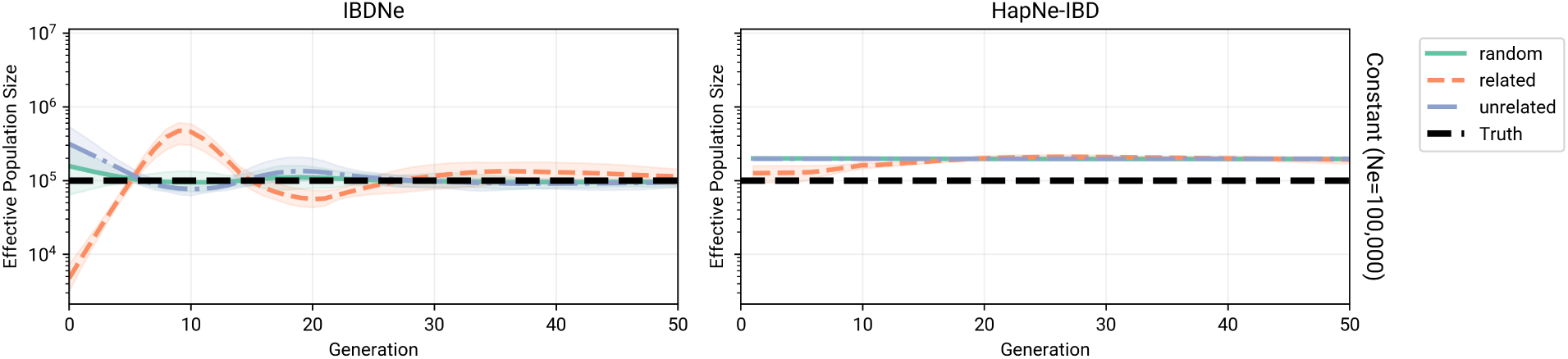
Inferred N*_e_*(t) for IBDNe and HapNe-IBD for a constant N*_e_* = 100, 000 population. Each line represents a different IBD filtering input into the model: random (green; individuals are selected at random), related (orange; 1st-3rd degree relatives are preferentially included), and unrelated (purple; 1st-3rd degree relatives are preferentially excluded). The mean N*_e_*(t) for 25 iterations is plotted and a 95% confidence interval is shaded. In general, in the “related” filtering recent N*_e_*(t) estimates are lower, particularly in IBDNe. HapNe-IBD’s smoothing procedure attenuates the choice of filtering, although it converges on an N*_e_*(t) roughly 2x the true N*_e_*(t). HapIBD suffers from oscillations going back as far as 30 generations in the past. The oscillations are more pronounced when relatives are oversampled (“related”) or undersampled (“unrelated“).s

## Notes

### Competing Interest Statement

The authors have declared no competing interest.

